# Changes in the expression of interleukin-10 in myocardial infarction and its relationship with macrophage activation and cell apoptosis

**DOI:** 10.1101/787887

**Authors:** Wenqi Yang, Shuming Li, Yang Zhao, Ying Sun, Yuling Huang, Zengli Diao, Cainai Xing, Fang Yang, Wenbo Liu, Xuan Zhao, Xiaoming Shang

**Affiliations:** Department of Hebei Medical University, Shijiazhuang, Hebei, United States of China; Department of North China University of Science and Technology, Caofeidian, Tangshan, Hebei, United States of China; Department of Hebei North University, Zhangjiakou, Hebei, United States of China; Department of Internal Medicine, North China University of Science and Technology Affiliated Hospital, Tangshan, Hebei, United States of China

**Keywords:** Myocardial infarction, IL-10, Macrophage, Cell apoptosis

## Abstract

Currently, the role of IL-10 as an anti-inflammatory factor in the occurrence and development of heart disease is still unclear. This study aimed to observe the dynamic changes in the expression of IL-10 in serum and myocardial tissues, as well as to investigate the relationship of IL-10 expression with macrophage activation and cardiomyocyte apoptosis during the occurrence of myocardial infarction. Mice models with myocardial infarction were prepared by ligating anterior descending branch of the coronary artery. The animals were classified into sham operation group (the control group), as well as groups of myocardial infarction based on days 1, 7, 14 and 28. On days 7 and 14, the cells with positive IL-10 expression were largely distributed in the infarct areas, while cells with positive IL-10 expression were decreased on day 28. Serum IL-10 was significantly positively correlated with IL-10 protein expression in myocardial tissues. Moreover, Bcl-2 and Bax protein expression in myocardial tissues, as well as the ratio of Bcl-2/Bax proteins were gradually elevated with prolonged time of infarction. There were positive correlations between IL-10 and Arginase expressions, and between the expressions of Bcl-2 and Bax proteins. After the occurrence of myocardial infarction, the expression of IL-10 was firstly increased and then decreased in serum and myocardial tissues, and this might affect macrophage activation, phenotypic transformation and the occurrence of cardiomyocyte apoptosis.

## Introduction

Coronary atherosclerosis, also known as coronary heart disease, is caused by myocardial ischemia, hypoxia and necrosis due to stenosis or complete occlusion of vascular lumen secondary to atherosclerosis of the coronary artery. Myocardial infarction is the most common type of heart disease [1,2]. Currently, most of the scholars hypothesized that immune and inflammatory responses are important causes for the onset of coronary heart diseases. Several types of inflammatory cells secrete and release various proinflammatory factors at the site of inflammation, and these are involved in the pathogenesis of coronary heart disease[2,3]. Myocardial cells undergo apoptosis in addition to ischemia and necrosis in patients with coronary heart diseases [4,5]. Currently, numerous apoptotic proteins are being studied, in which B cell lymphoma-2 (Bcl-2) protein family is the most commonly investigated and is the most classical apoptotic protein [5,6].

Recent studies have revealed interleukin-10 (IL-10) as an anti-inflammatory factor that plays an important role in the aspects of stability maintenance of coronary atherosclerotic plaque, tissue repair, etc., which also plays a “protective” role in the occurrence and development of coronary heart disease [7-9]. Most of the inflammatory cells (including macrophages) can synthesize and secrete IL-10^9^. Nevertheless, it is currently unclear as to whether IL-10 expression is internally correlated with macrophage activation and cardiomyoctyte apoptosis in coronary heart disease (including acute myocardial infarction).

## Materials and Methods

### Establishment of experimental animals and models with myocardial infarction

Thirty-one healthy male C57BL/6 mice of clean grade aged 16-20 weeks were fed in a clean animal room in the Medical Experimental Center of North China University of Science and Technology. The Animals were anesthetized with 1% pentobarbital sodium (50 mg/kg) via intraperitoneal injection, and cared in accordance with the principles of the Committee of the North China University of Science and Technology, which is consistent with the National Institutes of Healty (NIH) guidelines for the care and use of laboratory animals. The experimental procedure was approval by the Ethics Committee of North China University of Science and Technology, Tangshan, China(2014-020).

Preparation of mice models with myocardial infarction: The mice were anesthetized by intraperitoneal injection of 1% pentobarbital sodium, followed by tracheal intubation, and so a ventilator was connected. A 2 cm incision was made at the left sternal margin to expose the heart between the 3^rd^ and 4^th^ costa. The anterior descending branch of the left coronary artery was ligated approximately 2 mm below the left atrial appendage using the 8-0 prolene thread. After successful ligation, the anterior wall of the left ventricle was turned into pale due to rapid ischemia. The wounds were sutured in sequence.

Preparation of mice models for the sham group (the control group): After the mice were anesthetized, the thoracic cavity was opened to expose the heart and the anterior descending branch of the left coronary artery. Threading was performed without ligation, and then the wound was sutured.

A total of 4 mice died (2 mice due to pneumothorax and 2 mice due to hemorrhage in the thoracic cavity) during animal experiment. Two mice died (due to hemothorax) 24 hours after the operation. The survival rate of mice models was 81%.

### Experimental grouping and material collection

The mice models were divided into 5 groups (with 5 mice in each group) in the experiment, which included the control group (C), the group of mice with myocardial infarction for 1 day (1 d), the group of mice with myocardial infarction for 7 days (7 d), the group of mice with myocardial infarction for 14 days (14 d) and the group of mice with myocardial infarction for 28 days (28 d).

After the mice were sacrificed by anesthesia according to the specified time points, blood was collected from the eyeballs and stored in a refrigerator at 4°C for 2 h. The blood was centrifuged for collecting the supernatant and then stored in a refrigerator at −80°C for later use. The heart was extracted and the left ventricle was cut into two parts longitudinally. One part was fixed using 4% paraformaldehyde, while the other part was stored in a refrigerator at −80°C.

### Preparation of histological sections and staining

The heart tissues of the mice were fixed with 4% paraformaldehyde for 24 hours. The tissues were then soaked and embedded in paraffin after gradient dehydration using ethanol and deethanolization using xylene. The paraffin tissues were cut into 4 μm slices. Pathological changes were observed by routine hematoxylin & Eosin (HE) staining, and collagen deposition was observed by MASSON staining. Distributions of IL-10 and Arginase in myocardial tissues were observed by immunohistochemical staining. Cell apoptosis was observed under a fluorescence microscope after TUNEL staining (Millipore USA).

### Detection of serum IL-10 expression (ELISA)

The expression of IL-10 in venous blood of mice was detected by enzyme-linked immunosorbent assay (ELISA). The operations were conducted based on the kit (Andy gene) instructions.

### Detection of protein expressions in myocardial tissues (Western blot)

A total of 50 mg of myocardial tissues were weighed, and 1 mL RIPA lysate containing protease inhibitor was added. After that, centrifugation was performed to extract the supernatant protein. The protein standard curve was plotted by Bradford method. 15 μg of protein that needs to be tested was loaded, and electrophoresis and electrolytic membrane were then performed. Primary antibodies (IL-10, type I collagen, type III collagen, Tubulin, MCP-1, Arginase, iNOS, Bcl-2 and Bax. Dilution was performed at 1:1000) and secondary antibodies (HRP-labeled goat anti-rabbit or goat anti-mouse IgG. Dilution was performed at 1:5000) were added. Color development was conducted by ECL luminescence reagent. The bands were analyzed using Image J image processing software. The ratio of optical density of the target protein to the optical density of internal (Tubulin) protein was used as the relative expression of the protein.

### Statistical analysis

SPSS 20.0 software was used for statistical analysis. Normality was tested for quantitative data, and the data that met normal distribution were represented by mean ± standard deviation. The mean values of multiple groups were compared using one-way ANOVA, while pair-wise comparison was performed using least significance difference (LSD) test. *P*<0.05 was considered to be statistically significant.

## Results

### Pathomorphological changes in mice with myocardial infarction

On day 1 of myocardial infarction, severe degeneration and necrosis in the in the infarct areas of the left ventricle, accompanied by inflammatory cell infiltration were observed in the mice. On day 7 of myocardial infarction, there were more inflammatory cell infiltrates in addition to degenerated and necrotic cardiomyocytes, the hyperplastic collagen fibers (blue in Masson staining) were extended till the infarcted myocardial tissues, and then the infarcted myocardial tissues were separated. On day 14 of myocardial infarction, there were still degenerated and necrotic cardiomyocytes, accompanied by more inflammatory cell infiltrates. Moreover, collagen hyperplasia was more significant, separating the infarcted areas and the myocardial tissues in the adjacent areas into “islands”. On day 28 of myocardial infarction, the inflammatory cell infiltrates were declined when compared to those on day 14, and the tissues in the infarct areas were replaced by hyperplastic collagen fibers, while a small amount of residual cardiomyocytes were observed during that period (The results of Masson staining, and the results of HE staining were omitted in Fig 1).

**Fig 1.**
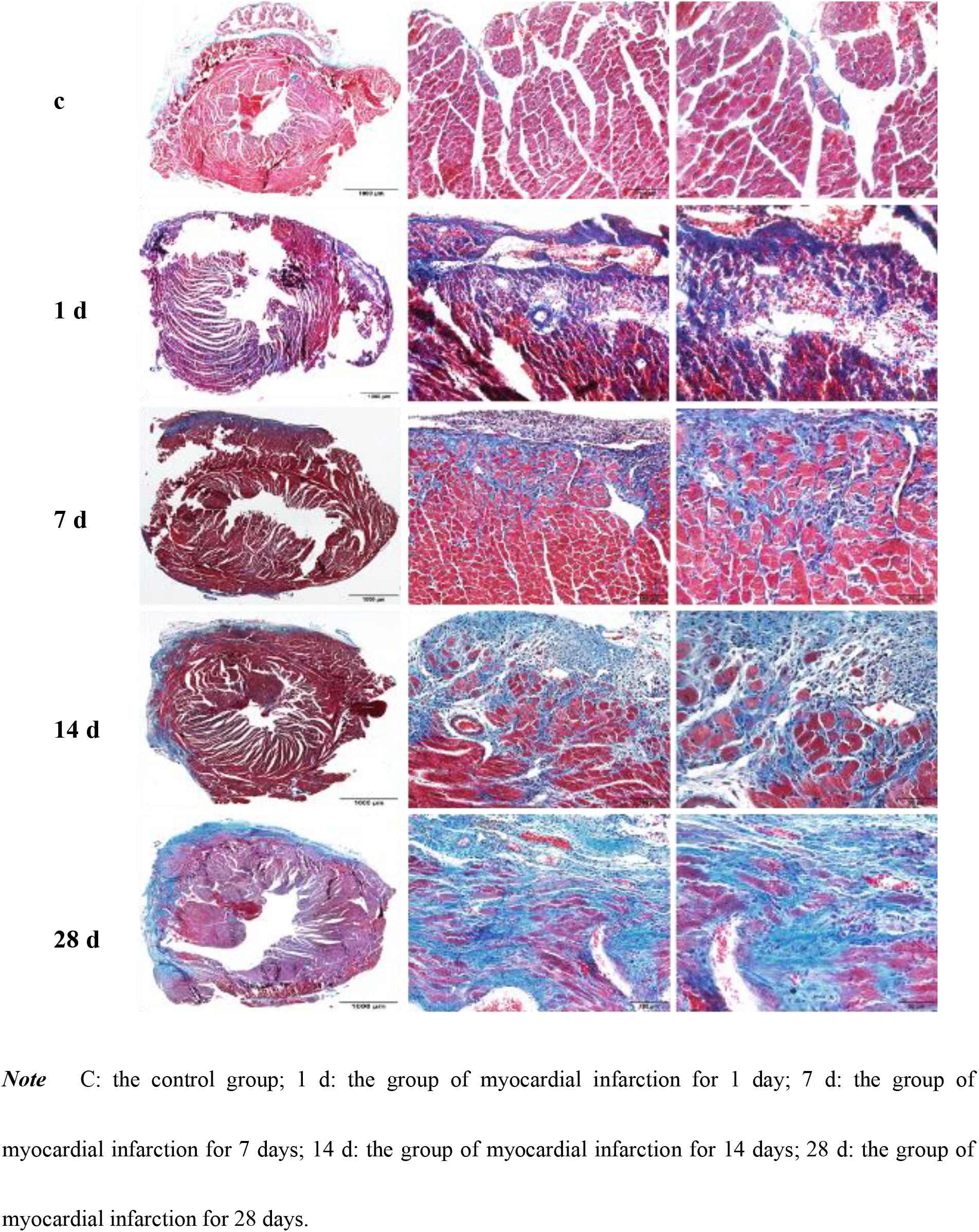
Pathological changes in mice with MI (Masson staining)

### Distribution and expression of IL-10 in the heart tissues of mice

Immunohistochemical staining demonstrated the existence of few IL-10 positive expression cells in the left ventricle in the control group. There were a small amount of cells with positive IL-10 expression in the infarcted areas on day 1 of myocardial infarction, there were a large number of cells with positive IL-10 expression on day 7 of myocardial infarction, and there were still a large number of cells with positive IL-10 expression on day 14, but cells with positive IL-10 expression was decreased on day 28. IL-10 was mainly expressed in the inflammatory cells in the lesion areas (Fig 2).

**Fig 2.**
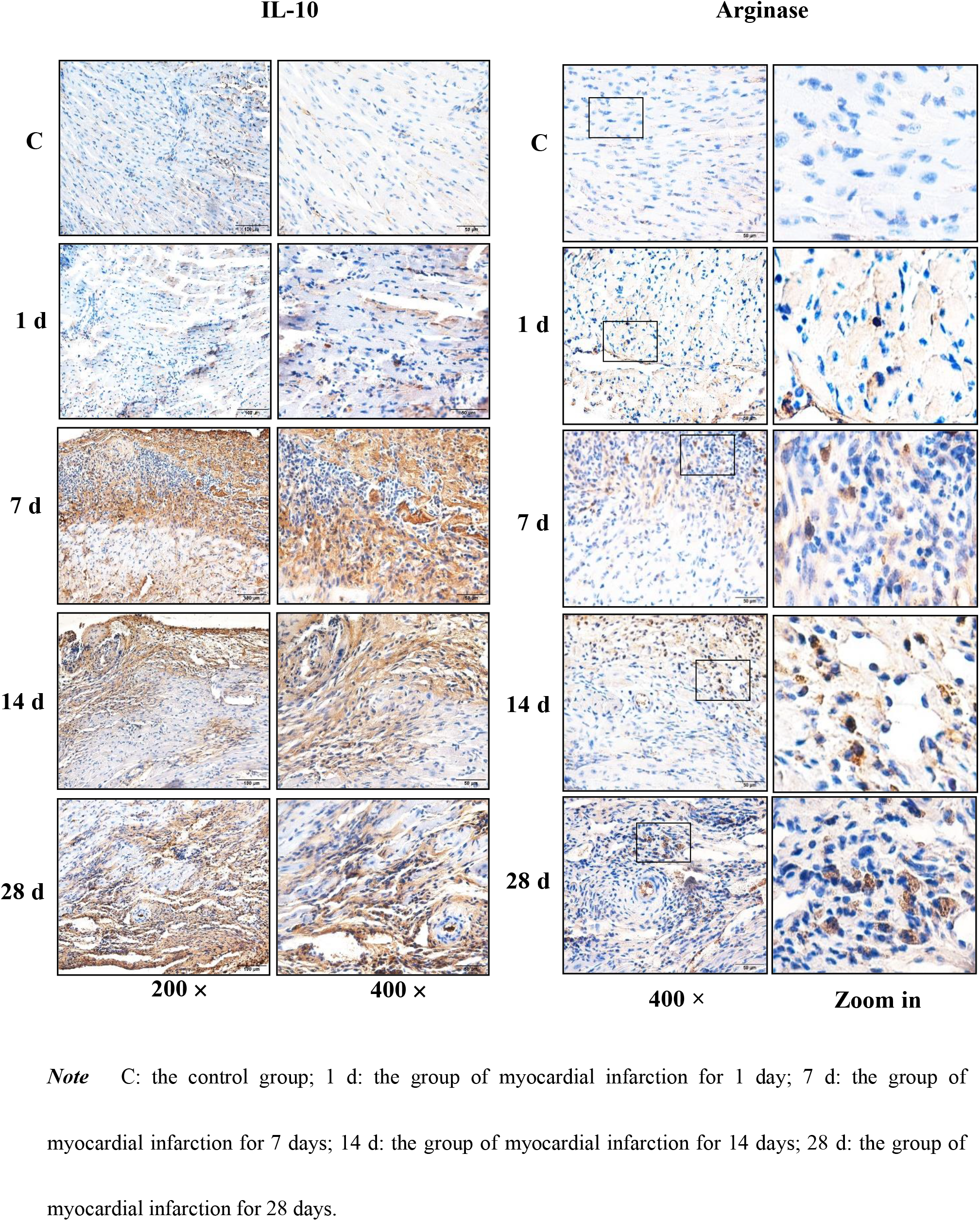
Distribution and change of IL-10 and Arginase in myocardial tissue of mice with MI (immunohistochemistry staining)

### Expression of serum IL-10

The results of ELISA demonstrated that the serum IL-10 expression was significantly increased when compared to the control group on days 1, 7 and 14 days of myocardial infarction, which were 1.19 times, 1.22 times and 1.18 times that of the control group, respectively, and the differences were statistically significant (*P*<0.05). However, on day 28 of infarction, the expression was decreased to a level approaching that of the control group (Table 1).

**Table 1.**
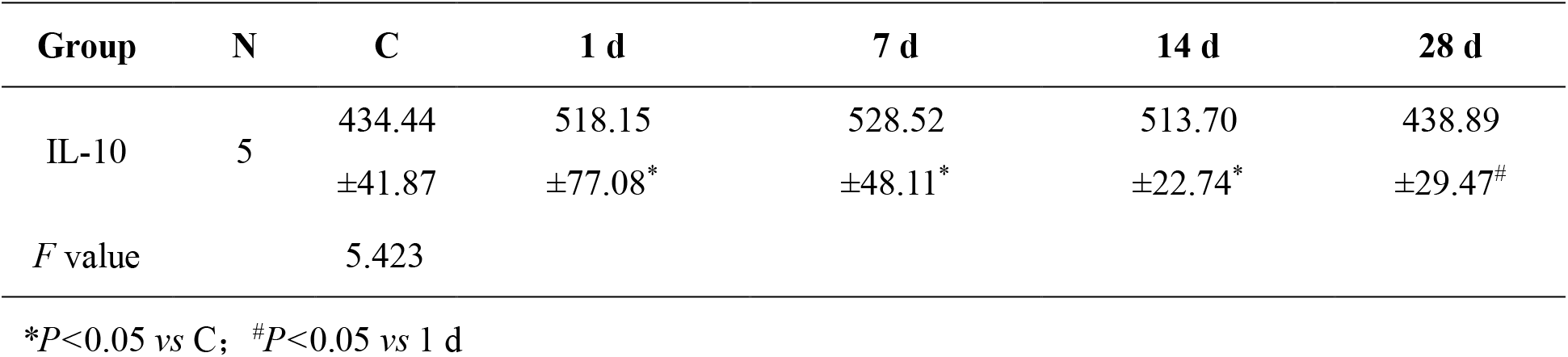
Changes of expression of IL-10 in serum of mice with MI during different time (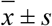, pg/ml, ELISA assay)

| Group | N | C | 1 d | 7 d | 14 d | 28 d |
| --- | --- | --- | --- | --- | --- | --- |
| IL-10 | 5 | 434.44<br>$\pm 41.87$ | 518.15<br>$\pm 77.08^*$ | 528.52<br>$\pm 48.11^*$ | 513.70<br>$\pm 22.74^*$ | 438.89<br>$\pm 29.47^\#$ |
| *F* value |  | 5.423 |  |  |  |  |

\* $P<0.05$ vs C; $^\#P<0.05$ vs 1 d

### Expression of IL-10 in myocardial tissues

The results of western blotting revealed that the expression of IL-10 protein in myocardial tissues was increased to a certain degree when compared to the control group on day 1 of myocardial infarction, and was significantly increased on day 7 of myocardial infarction, which was 1.43 times that of the control group (*P*<0.05). Moreover, IL-10 expression was peaked on day 14 of myocardial infarction, which was 1.7 times that of the control group. However, on day 28 of myocardial infarction, the expression of IL-10 protein was decreased significantly when compared to that on day 14, which was 74.48% that of day 14 (*P*<0.05, Fig 3).

**Fig 3.**
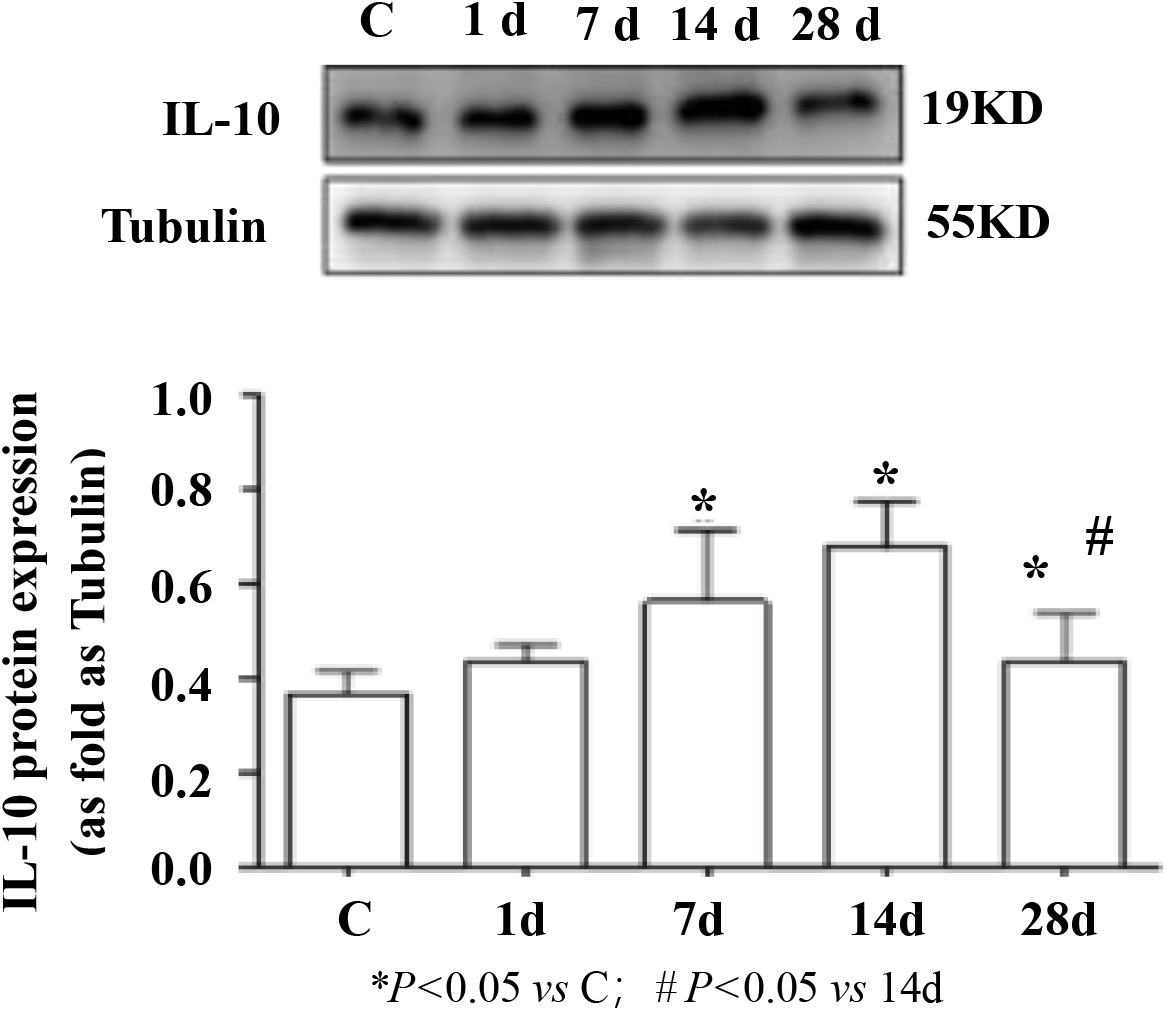
Changes of myocardial IL-10 protein expression in mice with MI (western blot)

Correlation analysis of serum IL-10 and IL-10 expression in myocardial tissues showed that r was 0.543 and P was 0.005, demonstrating that the two were positively correlated.

### Distribution and changes of M2 macrophages in the infarcted areas of mice with myocardial infarction

Immunohistochemical staining results showed that there were no cells with significantly positive expression of Arginase (that is M2 macrophages) distributed in the myocardial tissues in the control group. On day 1 of myocardial infarction, the cells with positive expression of Arginase were occasionally visible in the infarcted areas, there were some cells with positive expression of Arginase on day 7, and there were more cells with positive expression of Arginase on day 14 when compared to those on day 7, while there were still many cells with positive expression of Arginase on day 28 of myocardial infarction (Figure 2).

### Changes in the expressions of MCP-1, iNOS and Arginase in mice with myocardial infarction

The results of western blotting revealed that there was an increasing trend first and then a decreasing expression of MCP-1 and iNOS proteins in the myocardial tissues. Compared to the control group, the expressions of MCP-1 protein were 1.08 times, 2.73 times, 2.29 times and 1.39 times that of the control group on 1, 7, 14 and 28 days of myocardial infarction, respectively, while the expressions of iNOS protein were 1.5 times, 4.48 times, 3.28 times and 1.43 times higher, respectively. However, there was a gradual increasing trend in the Arginase protein expression. Compared to the control group, the expression of Arginase protein was 1.54 times, 2.42 times, 3.78 times and 4.12 times that of the control group on days 1, 7, 14 and 28 of myocardial infarction, respectively (Fig 4).

**Fig 4.**
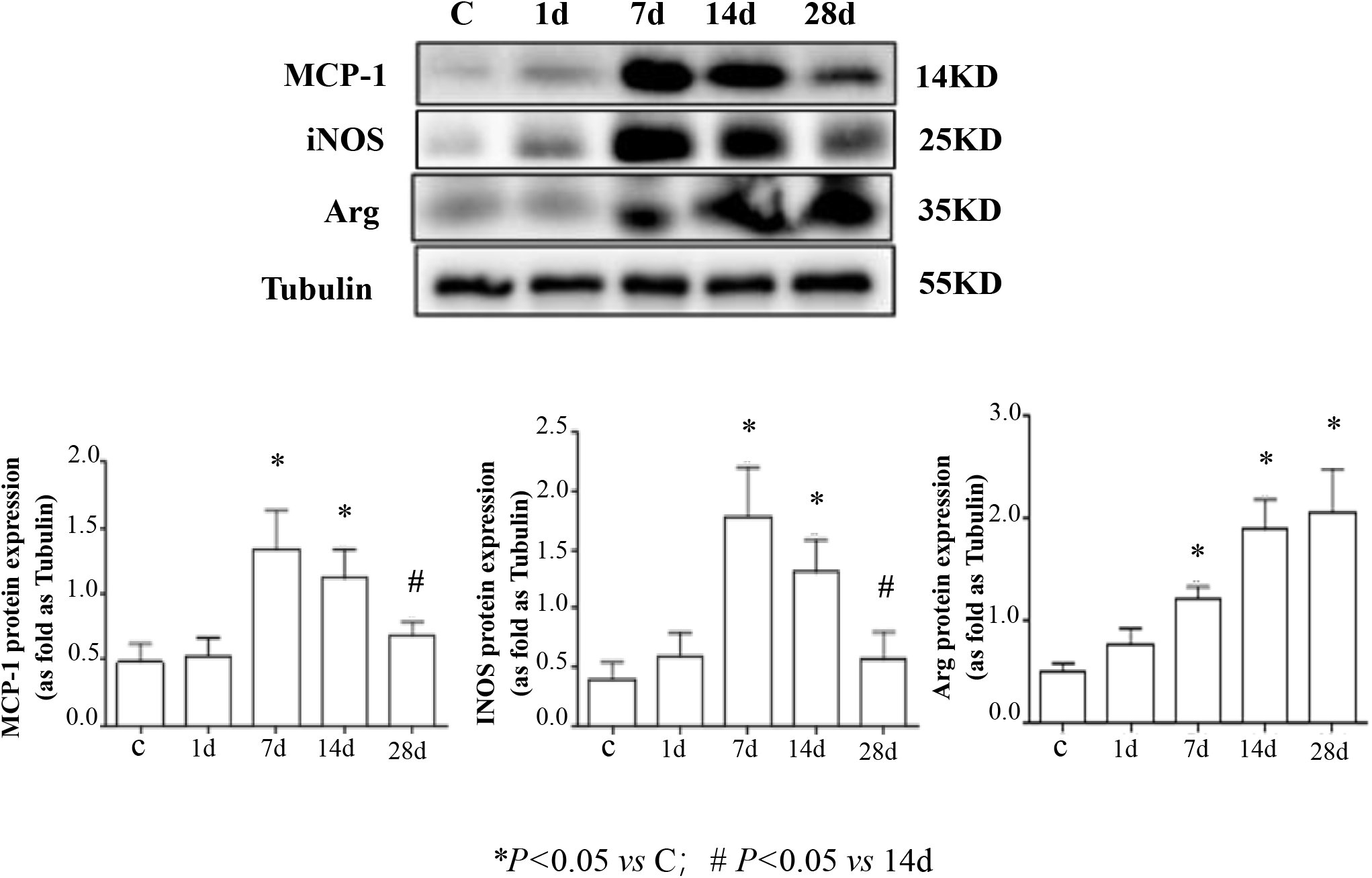
Changes of protein expression of MCP-1, iNOS and Arginase in myocardial tissue of mice with MI (western blot)

### Cardiomyocyte apoptosis of mice with myocardial infarction

The results of Tunel staining showed that individual apoptotic cells were occasionally visible in the myocardial tissues in the control group. On day 1 of myocardial infarction, the number of apoptotic cells in the infarcted areas was increased to some extent, and the number of apoptotic cells was increased gradually with prolonged time of infarction (Figure 5).

**Fig 5.**
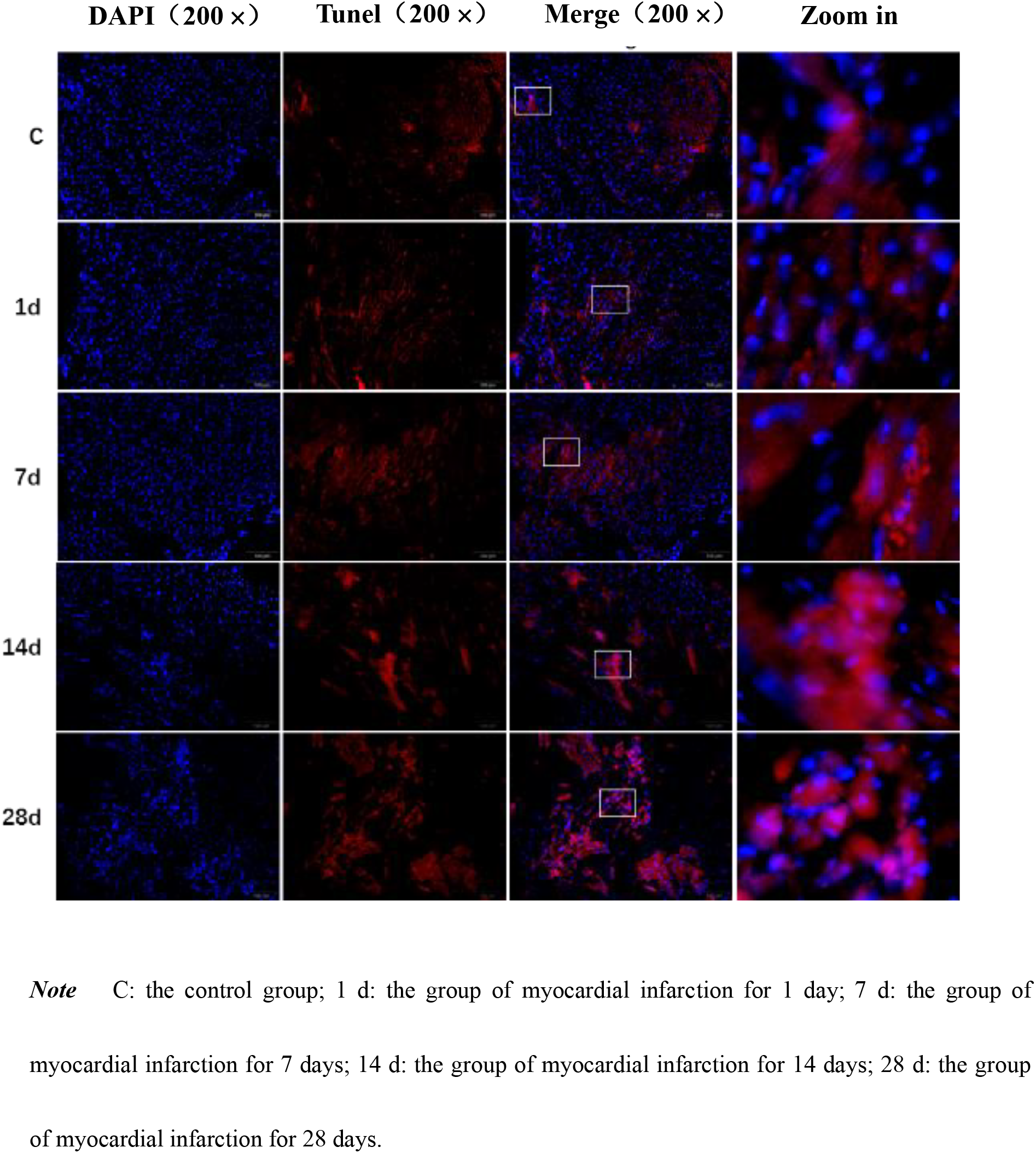
Morphology changes of apoptosis in mice with MI (Tunel staining)

### Changes in the expressions of Bax and Bcl-2 proteins in the myocardial tissues of mice with myocardial infarction

Western blotting revealed the expressions of Bcl-2 and Bax proteins, as well as the ratio of Bcl-2/Bax proteins was gradually increased with prolonged time of infarction. Compared with the control group, the expressions of Bax protein on days 1, 7, 14 and 28 of myocardial infarction were 1.19 times, 1.50 times, 1.70 times and 2.23 times those of the control group, while the expressions of Bcl-2 protein were 1.44 times, 3.08 times, 4.54 times and 6.04 times (Fig 6). The ratios of Bcl-2/Bax were 1.22 times, 2.03 times, 2.67 times and 2.74 times than those of the control group (Fig 7).

**Fig 6.**
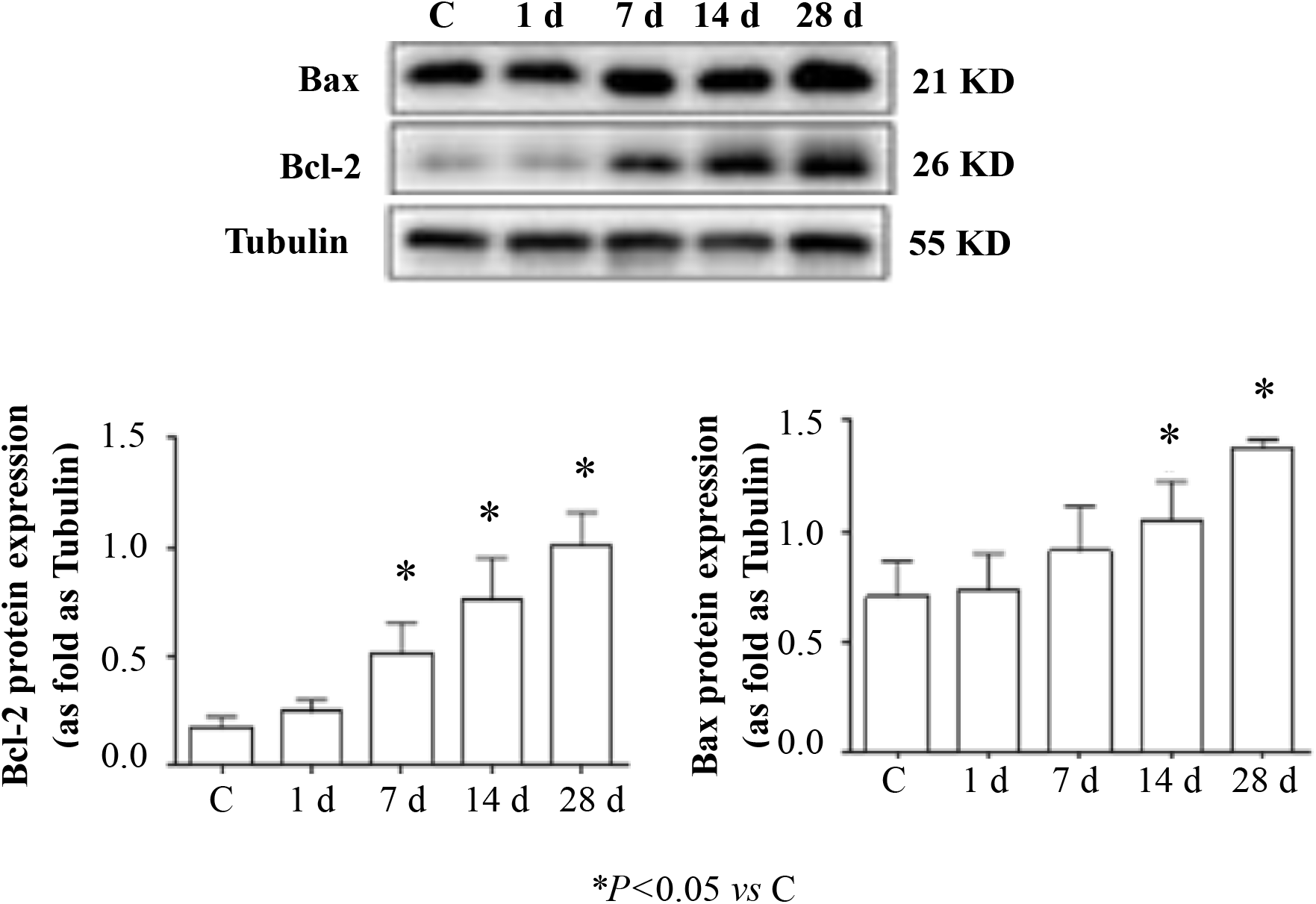
Changes of myocardial protein expression of Bcl-2 and Bax in mice with MI (western blot)

**Fig 7.**
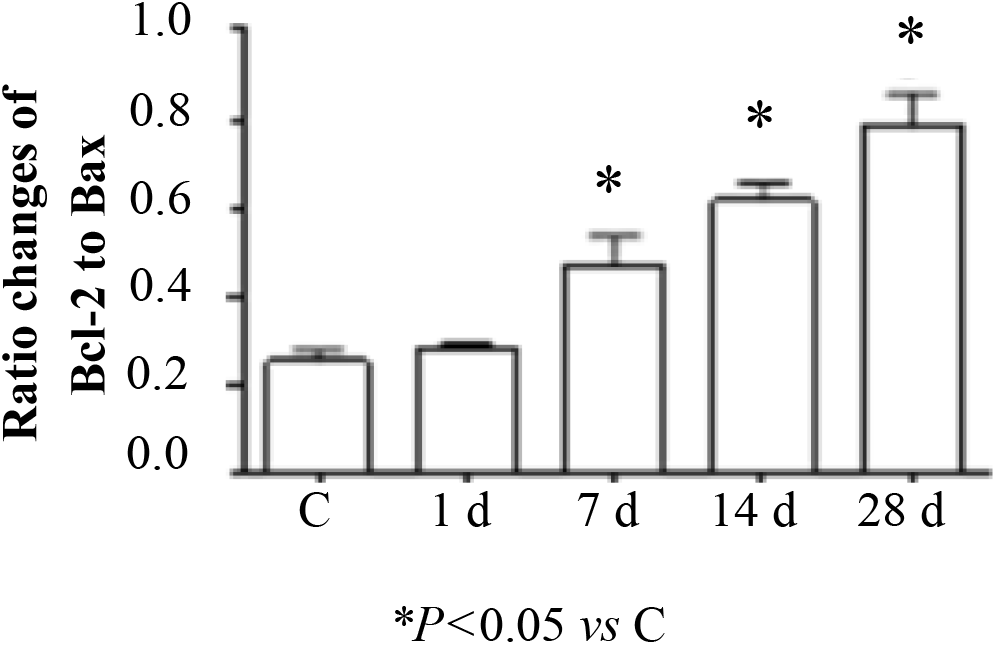
The ratio changes of Bcl-2 to Bax in myocardial protein expression of mice with MI (western blot)

### Correlation analysis of various proteins in myocardial tissues

Correlation analysis suggested a positive correlation among IL-10, Bcl-2, Bax and Arginase protein expressions (Table 2).

**Table 2.** Correlation analysis between several protein expressions in myocardial tissue

|  | IL-10 |  | Bcl-2 |  | Bax |  | Bcl-2/ Bax |  | Arginase |  |
| --- | --- | --- | --- | --- | --- | --- | --- | --- | --- | --- |
|  | r | P | r | P | r | P | r | P | r | P |
| IL-10 | — |  | 0.690 | 0.004 | 0.546 | 0.035 | 0.588 | 0.021 | 0.669 | 0.006 |
| Bcl-2 | 0.690 | 0.004 | — |  | 0.901 | 0.000 | 0.951 | 0.000 | 0.965 | 0.000 |
| Bax | 0.546 | 0.035 | 0.901 | 0.000 | — |  | 0.912 | 0.000 | 0.973 | 0.000 |
| Bcl-2/Bax | 0.588 | 0.021 | 0.951 | 0.000 | 0.912 | 0.000 | — |  | 0.951 | 0.000 |
| Arginase | 0.669 | 0.006 | 0.956 | 0.000 | 0.973 | 0.000 | 0.951 | 0.000 | — |  |

## Discussion

The inflammatory responses after myocardial infarction act as an important pathological process in myocardial infarction. A large number of inflammatory factors with different functions play an important role in the inflammatory responses [2,10]. IL-10 is an important inflammatory factor, which play a strong anti-inflammatory role. Studies have demonstrated that IL-10 is closely related with the occurrence and prognosis of cardiovascular diseases, and has important regulatory effect on remodeling of tissue functions and structures after ischemic injury[8,9,11]. Recently, it has been reported that the target cells in which IL-10 plays anti-inflammatory effects are mainly macrophages. Mononuclear macrophages secrete IL-10 after activation by various endogenous and exogenous mediators, causing IL-10 genetic transcription via lipopolysaccharide activation pathway, protein kinase A-dependent activation pathway, etc[12,13]. Additionally, mononuclear macrophages also secrete IL-10 during the process of clearing apoptotic cells, and this process depends on the activation of p38 mitogen-activated protein kinase (MAPK) [14]. It has been reported that the expression of IL-10 is increased during ischemia, which acts as a protective response to ischemia [15,16]. Some scholars have found that serum Il-10 significantly increases on the day of acute myocardial infarction. Moreover, IL-10 remains at a higher level on day 1 after percutaneous coronary intervention (PCI) therapy, but it decreases to some extent on day 7 after therapy, suggesting that IL-10 is associated with time and severity of myocardial ischemia to some extent [17]. Our study results showed that serum IL-10 expression was increased significantly on day 1 of myocardial infarction in mice, which remained at a higher level on the days 7 and 14 of myocardial infarction, but was decreased after that, and it decreased to a level that was close to that of the control group on day 28. The existence of a small number of cells with positive IL-10 expression in the infarcted areas on day 1 of myocardial infarction. Moreover, majority of the cells with positive IL-10 expression were visible on days 7 and 14 of myocardial infarction, while the cells with positive IL-10 expression were decreased on day 28. However, the expression of myocardial IL-10 protein was significantly elevated on day 7 of myocardial infarction, and reached peaked on day 14, while it was decreased significantly on day 28. Correlation analysis showed that the expression of serum IL-10 was positively correlated with the expression of IL-10 protein in myocardial tissues. These results suggested an increasing trend initially and then a decrease in IL-10 expression (regardless of in serum or myocardial tissues) with prolonged time of infarction in the mice models with myocardial infarction. On day 1, myocardial infarction was considered as the initial stage of acute inflammation, thus more necrosis of cardiomyoctyes, accompanied by exudation and infiltrates of inflammatory cells were observed. However, day 7 of myocardial infarction was the critical stage of acute inflammation, in which a large amount of exudation and infiltrates of inflammatory cells, accompanied by the release of a large number of proinflammatory factors were observed. The body in response to these releases anti-inflammatory factors (such as IL-10) to maintain a balance in the biological functions [11,18]. Therefore, at around days 1 and 7 of myocardial infarction, IL-10 was highly expressed in serum and/or myocardial tissues, which continued until week 2. After that, tissue injury entered the repair stage, inflammation was degraded gradually as well, and there was more extracellular matrix (including collagen) to replace the infarcted myocardial tissues gradually [3,19]. At that time, synthesis and secretion of IL-10 was gradually decreased, suggesting that the dynamic changes of IL-10 in serum and myocardial tissues during the process of myocardial infarction were closely associated with the occurrence and development of inflammation, as well as the outcomes of myocardial infarction. These might be caused due to the defense and anti-inflammatory effects of the body to inflammatory responses during myocardial infarction.

In inflammatory responses, macrophages are activated and phenotypic transformation occurs with the progression of inflammation. In the initial and early stages of inflammation, macrophages act as "proinflammatory" macrophages (M1 phenotype, secreting specific marker proteins such as TNF-α, iNOS and IL-1β, etc.), which in turn promote the progression of inflammation. However, in the middle and advanced stages, macrophages act as “repairing” macrophages (M2 phenotype, secreting specific marker proteins such as Arginase and CD206, etc.), playing their effects as anti-inflammation and tissue repair [19,20]. In addition, in vitro experiment showed that fibroblasts were activated after IL-10 intervention. Moreover, its proliferation and migratory abilities were strengthened, and there were more collagens that are closely related to M2 phenotypic transformation of macrophages [20]. In this study, degeneration and necrosis of cardiomyocytes were observed on day 1 of myocardial infarction in mice, the expression of inflammatory factor monocyte chemoattractant protein-1 (MCP-1) began to increase, and there was inflammatory cell infiltrates in the infarcted areas. Results of immunohistochemical staining showed the existence of only a small number of M2 macrophages in these inflammatory cells. Meanwhile, iNOS that represent the marker protein of MI macrophages was increased to some extent. On day 7 of myocardial infarction, inflammatory cell infiltration in the infarcted areas was more significant. Among them, the expression of MCP-1 protein showed a peak, and iNOS expression achieved the highest value. However, the number of cells (M2 macrophages) with positive Arginase expression was increased significantly, while the expression of Arginase protein was also elevated significantly. On day 14 of myocardial infarction, the iNOS expression showed declination with decreasing expression of MCP-1 protein. Until day 28, the expressions of MCP-1 and iNOS proteins were dropped back to the levels as in the control group, while the number of cells with positive Arginase expression still remained more, and the expression of Arginase protein was still elevated. This suggested that the macrophages were activated and there was a phenotypic transformation from M1 macrophages in the initial stage of inflammation to M2 macrophages in the middle and advanced stages of inflammation during the process of myocardial infarction. Correlation analysis showed that IL-10 expression in myocardial tissues was positively correlated with the expression of Arginase protein during the occurrence and development of myocardial infarction. This indicated that the dynamic changes of IL-10 after myocardial infarction might affect or regulate macrophage activation, and promote the transformation of proinflammatory macrophages to repair-type macrophages.

Cardiomyocyte apoptosis also occurs with the degeneration and necrosis of cells during the process of myocardial infarction. Bax and Bcl-2 are important cell apoptosis-related genes and cell apoptosis regulatory proteins. Bax plays a positive regulatory effect during cell apoptosis, promoting the occurrence of cell apoptosis. Bcl-2 plays a negative regulatory role, blocking the apoptotic pathway and inhibiting the occurrence of apoptosis [5,21]. However, the ratio of Bcl-2/Bax is closely related to cell apoptosis and survival [6,22]. In this study, the results showed that the number of apoptotic cells in myocardial tissues was increased gradually with prolonged time of infarction, and the expression of Bax protein was also increased gradually, indicating that cardiomyoctyte apoptosis was gradually aggravated with prolonged time of infarction. With prolonged time of infarction, the expression of Bcl-2 protein was elevated gradually, especially the ratio of Bcl-2/Bax was gradually increased. Correlation analysis showed that the expression of IL-10 protein in myocardial tissues was significantly positively correlated with the expressions of Bax and Bcl-2 proteins, especially the ratio of Bcl-2/Bax proteins. These dynamic changes suggested that IL-10 expression would affect the occurrence of cardiomyocyte apoptosis during the occurrence and development of myocardial infarction. Due to the gradual increase in the ratio of Bcl-2 to Bax, the effects of IL-10 might regulate cardiomyocyte apoptosis by controlling the balance between anti-apoptotic gene (Bcl-2) and apoptotic gene (Bax), thereby affecting remodeling of functions and structures of myocardial tissues.

In addition, further correlation analysis showed a positive correlation between IL-10 and Arginase in myocardial tissues as well as between the expressions of Bcl-2 and Bax proteins. This suggested that IL-10, Arginase and apoptosis-regulatory proteins showed close association, playing an important role in the remodeling of tissue structures and damage repair process after myocardial infarction.

## Acknowledgments

The authors would like to thank collaborators at the School of Public Health, North China University of Science and Technology. A special thanks goes to Miss Sun Ying, head of laboratory of Yang fang and his team who conducts IL-10 analyses.

